# Comparative dynamics of gene expression during in vitro and in vivo *Candida albicans* filamentation

**DOI:** 10.1101/2023.09.21.558874

**Authors:** Rohan S. Wakade, Damian J. Krysan

## Abstract

*Candida albicans* is one of them most common causes of fungal disease in humans and is a commensal member of the human microbiome. The ability of *C. albicans* to cause disease is tightly correlated with its ability to undergo a morphological transition from budding yeast to a filamentous form (hyphae and pseudohyphae). This morphological transition is accompanied by the induction of a set of well characterized hyphae-associated genes and transcriptional regulators. To date, the vast majority of data regarding this process has been based on in vitro studies of filamentation using a range of inducing conditions. Recently, we developed an in vivo imaging approach that allows the direct characterization of morphological transition during mammalian infection. Here, we couple this imaging assay with in vivo expression profiling to characterize the time course of in vivo filamentation and the accompanying changes in gene expression. We also compare in vivo observations to in vitro filamentation using a medium (RPMI 1640 tissue culture medium with 10% bovine calf serum) widely used to mimic host conditions. From these data, we make the following conclusions regarding in vivo and in vitro filamentation. First, the transcriptional programs regulating filamentation are rapidly induced in vitro and in vivo. Second, the tempo of filamentation in vivo is prolonged relative to in vitro filamentation and the period of high expression of genes associated with that process is also prolonged. Third, hyphae are adapting to changing infection environments after filamentation has reached steady-state.

**Importance:** *Candida albicans* filamentation is correlated with virulence and is an intensively studied aspect of *C. albicans* biology. The vast majority of studies on *C. albicans* filamentation are based on in vitro induction of hyphae and pseudohyphae. Here we used an in vivo filamentation assay and in vivo expression profiling to compare the tempo of morphogenesis and gene expression between in vitro and in vivo filamentation. Although the hyphal gene expression profile is induced rapidly in both conditions, it remains stably expressed over the 24hr time course in vivo while the expression of other environmentally responsive genes is dynamic. As such, it is important to regard the filamentation process as a separate growth phase of *C. albicans* that is as adaptable to changing growth conditions as the more familiar yeast phase.

## Introduction

*Candida albicans* is a component of the human mycobiome that also causes disease in both immunocompetent and immunocompromised patients (1, 2). The transition of *C. albicans* from harmless commensal to invasive pathogen is associated with a morphological switch from budding yeast to filaments comprised of both hyphae and pseudohyphae (3). In general, *C. albicans* strains and mutants with reduced filamentation are more fit in the commensal setting and less fit during invasive infection (4, 5). Over the years, filamentation has been one of the most intensively studied *C. albicans* virulence traits (3). The vast majority of these studies have focused on filaments generated in vitro using a wide range of filament inducing conditions. Recently, we developed an in vivo imaging approach to the analysis of *C. albicans* filamentation during infection that is based on the direct inoculation of the subdermal ear tissue of the mouse (6). We have used this approach to identify the transcription factor network that regulates filamentation in vivo (7) as well as to characterize the filamentation of clinical isolates and protein kinase mutants (8). These studies have revealed a number of important differences in the genes required for filamentation in vivo compared to in vitro. For example, the cAMP-Protein Kinase A pathway is absolutely required for in vitro filamentation in all inducing-media but is dispensable for filamentation in vivo (9).

As part of these studies, we have also used Nanostring to characterize the in vivo expression of a set of 186 environmentally responsive genes (7, 9). This set of genes includes 57 (30%) hyphae-associated genes (7). The Nanostring-based in vivo expression profile of *C. albicans* in ear tissue to infection of the kidney and to in vitro-induced filaments showed a reasonable correlation but also indicated the existence of differences (9). To further explore the differences in the genetic regulation of filamentation between in vivo and in vitro conditions, we followed the time course of both morphogenesis and gene expression during in vitro and in vivo filamentation. We used the Nanostring based approach for two primary reasons. First, genome-wide RNA-seq methods for the direct analysis of *C. albicans* gene expression in infected tissue have not been developed. Although a gene-enrichment strategy has been reported (10), its application to time course analysis was cost prohibitive. Second, we were most interested in the temporal dynamics of a set of well-studied hyphae-associated genes and genes that are responsive to environmental conditions in *C. albicans* (11). Although this approach limits the conclusions we can make about global patterns of gene expression, the patterns observed provide insights into important aspects of the morphogenesis process and the distinctions in the regulation of specific, reporter genes between in vitro and in vivo morphogenesis.

We used a single in vitro filament-inducing condition which is generally considered to be “host-like” in the field. Specifically, the tissue culture medium RPMI 1640 supplemented with 10% bovine calf serum (BCS) was used and incubations were performed at mammalian body temperature (37°C). Previous microarray-based expression profiling of the time course for in vitro *C. albicans* filamentation used rich medium (YPD or YEPD) supplemented with 10% BCS (12). It has become well-established that the specific filament-inducing medium has a significant effect on both the regulatory pathways and gene expression patterns involved in *C. albicans* filamentation (13). Indeed, those considerations motivated our study of gene expression pattern in vivo.

With these limitations in mind, our data show that the vast majority of hypha-associated genes are rapidly induced both in vitro and in vivo. In vivo, a steady state ratio of filaments to yeast is reached between 8 and 12 hr post-infection and is maintained to 24 hr with sustained expression of hyphae-associated genes. In vitro, the expression of hyphae-associated genes begins to fall by 4 hr, a point where lateral yeast formation begins to occur (14). We also show that gene expression patterns continue to change during the morphologic steady state in vivo with metabolic and cell wall genes undergoing significant changes.

## Results

### In vivo filamentation reaches steady state twelve hours after infection

We previously characterized *C. albicans* filamentation 24 hr after inoculation of the ear tissue (7, 9). This time point was used partly because the ratio of filaments to yeast at that time point was similar to that observed after 4 hr of induction with RPMI 1640 supplemented with 10% bovine calf serum (our standard vitro conditions unless otherwise indicated). However, we had not performed a formal time course analysis to characterize the dynamics of morphology in the model. Unfortunately, the initial inflammation of inoculation prevented consistent analysis of filamentation prior to the 8 hr time point. We were, however, able to measure both the ratio of filaments to yeast and the length of filaments at 8 hr and 12 hr and compare it to our previous data at 24 hr. The ratio of filaments to yeast is ∼ 20% lower at 8 hr compared to either 12 hr or 24 hr (Fig. 1A) but there is no significant difference between the 12 hr and 24 hr time points. Although the difference in filament length is also reduced at 8 hr compared to 24 hr, the absolute difference in length is modest (∼3 mm, Fig. 1B); the difference between 12hr and 24hr is not significant. This indicates that a morphological steady state is established between 8 and 12 hr after which the ratio of filament to yeast remains fairly stable.

**Figure 1.**
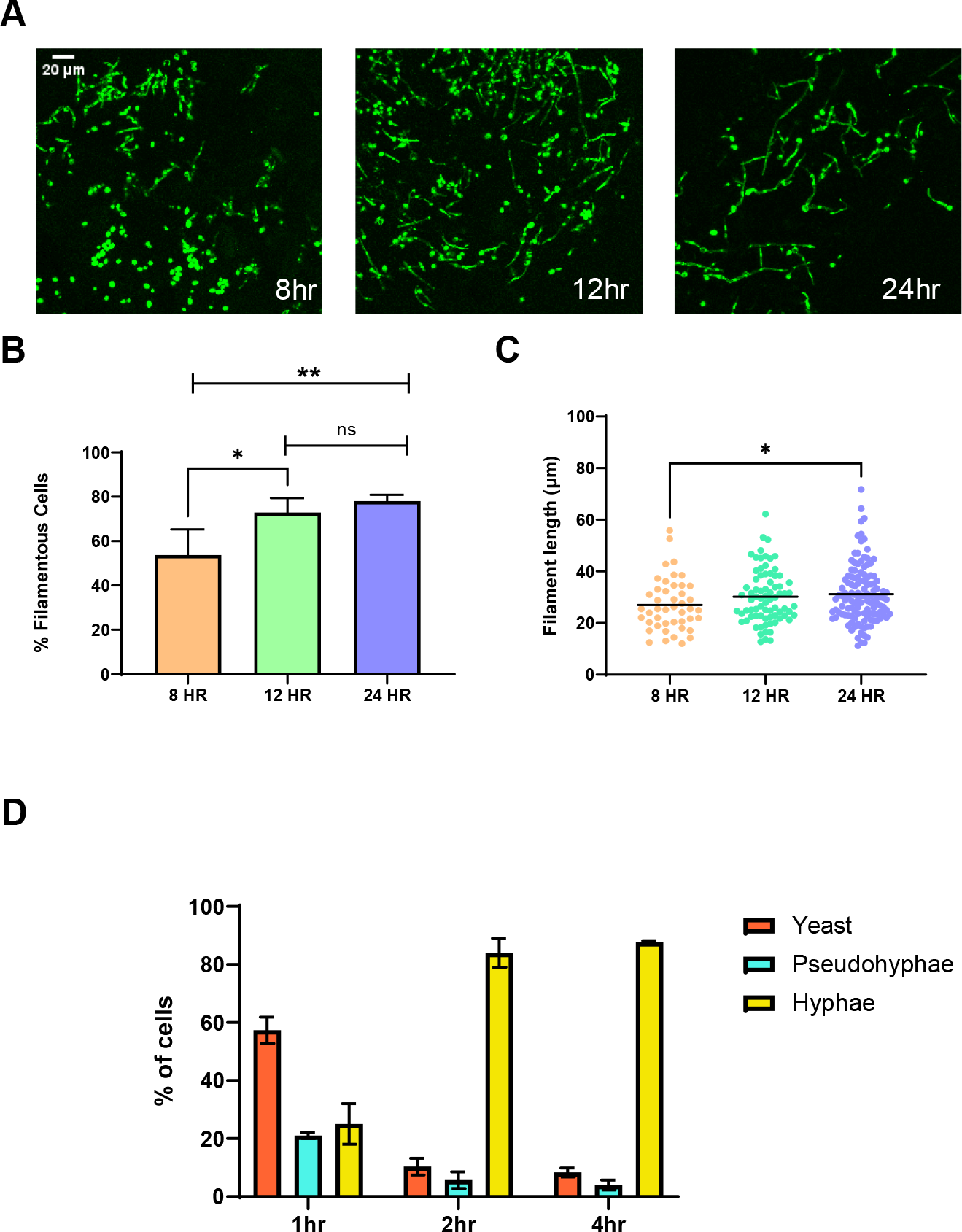
Time course of in vivo and in vitro *C. albicans* filamentation. **A.** Representative 2-D fields from confocal microscopy of NEON-labeled SN250 strain in ear tissue at the indicated time points after infection. **B**. Quantification of percent filamentous cells at the indicated time points. Asterisks indicated significant difference between time points (p < 0.05 *, p <0.005 **, Student’s t test); NS indicates no significant difference. **C**. The length of filaments at the indicated time points. Asterisks indicate significant difference between the groups (p < 0.05 *, Mann Whitney test). **D**. Distribution of hyphae, pseudohyphae and yeast for cells inoculated into RPMI + 10% BCS at 37°C for the indicated periods of time.

We performed a more condensed time course for in vitro induction with time points at 1hr, 2hr, and 4hr (Fig. 1D). At 1 hr post-inoculation, the cultures remain predominantly yeast form. At 2hr, ∼80% of the cells are hyphae and that proportion remained the same at 4 hr. The extent of filamentation at the 4hr time point was similar to that at 12 or 24hr in vivo. After 4hr incubation in filament inducing media, lateral yeast forms begin to emerge from the hyphal filaments with approximately 10% of hyphae containing lateral yeast; as others have also shown, the extent of lateral yeast formation then increases over time (14, 15). In vivo, however, we have not previously observed lateral yeast formation at the 24hr time point and we did not observe lateral yeast at any of the time points in this work (7, 8, 9). These data indicate that in vivo filamentation occurs at a reduced rate relative to in vitro inductions and that a steady state of morphotypes is established between 8 and 12 hours after inoculation.

### Similarity and differences in the expression of morphogenesis reporter genes in vitro and in vivo

We used Nanostring nCounter methods to characterize the expression of 186 genes and compared them to the yeast inoculum at 1, 2, 4, 8, and 12hr post-infection (Table S1) as well as 1 and 4hr post filament induction in vitro (Table S2). These data were also compared to those previously collected at 24hr using the same *C. albicans* and mouse strains (8,9) as well as to 4 hr previously generated using the same medium and strain. The raw, normalized, and processed data are provided in Table S1/2 for each time point along with previously published data for 24hr in vivo (9). The time course of transcriptional changes is summarized by volcano plots in Fig. 2A and 2B for in vivo and in vitro conditions, respectively. The total number of differentially expressed genes at each time point is indicated in Fig. 2A and 2B.

**Figure 2.**
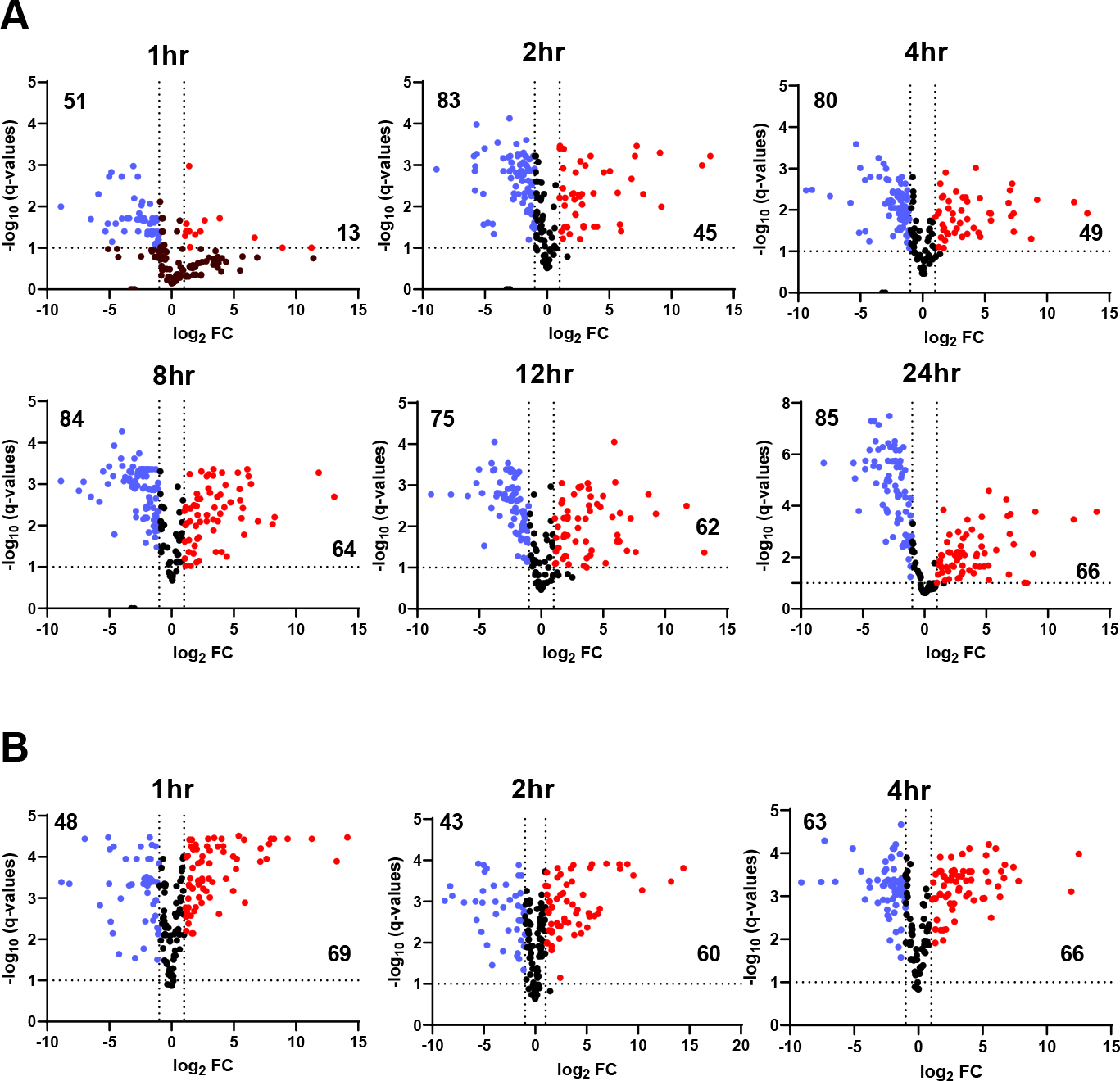
Nanostring analysis of gene expression over time for in vivo and in vitro filamentation. Volcano plots showing genes with significant (log_2_ ± 1; FDR<0.1, Benjamini method) increase (red dots) or decrease (blue dots) at the indicated time points for in vivo filamentation (**A**) and in vitro filamentation (**B**). Expression is normalized to yeast phase cells used to infect mice or inoculate in vitro cultures. The numbers in the two quadrants indicate total number of differentially expressed genes.

We first focused on the expression of hyphae-specific genes and hyphae-induced transcriptional regulators over the in vivo and in vitro time courses. *HWP1* is a widely used reporter of the hyphal transcriptional program (16) while *YWP1* is expressed in yeast and downregulated in hyphae (17). The time course for the expression of these two genes is shown in Fig. 3A&B for in vivo and in vitro conditions, respectively. *HWP1* expression is rapidly induced relative to the yeast phase inoculum both in vivo and in vitro. The expression reaches a plateau in vivo between 2 and 4 hrs and then is stable for the remainder of the time course. In vitro the plateau is reached after 1 hr. *YWP1* expression declines more slowly in vivo and reaches a steady state of expression at 8 hr. Relative to the time course of filamentation under in vitro and in vivo conditions, both *HWP1* and *YWP1* expression have reached steady state before morphological steady state is reached.

**Figure 3.**
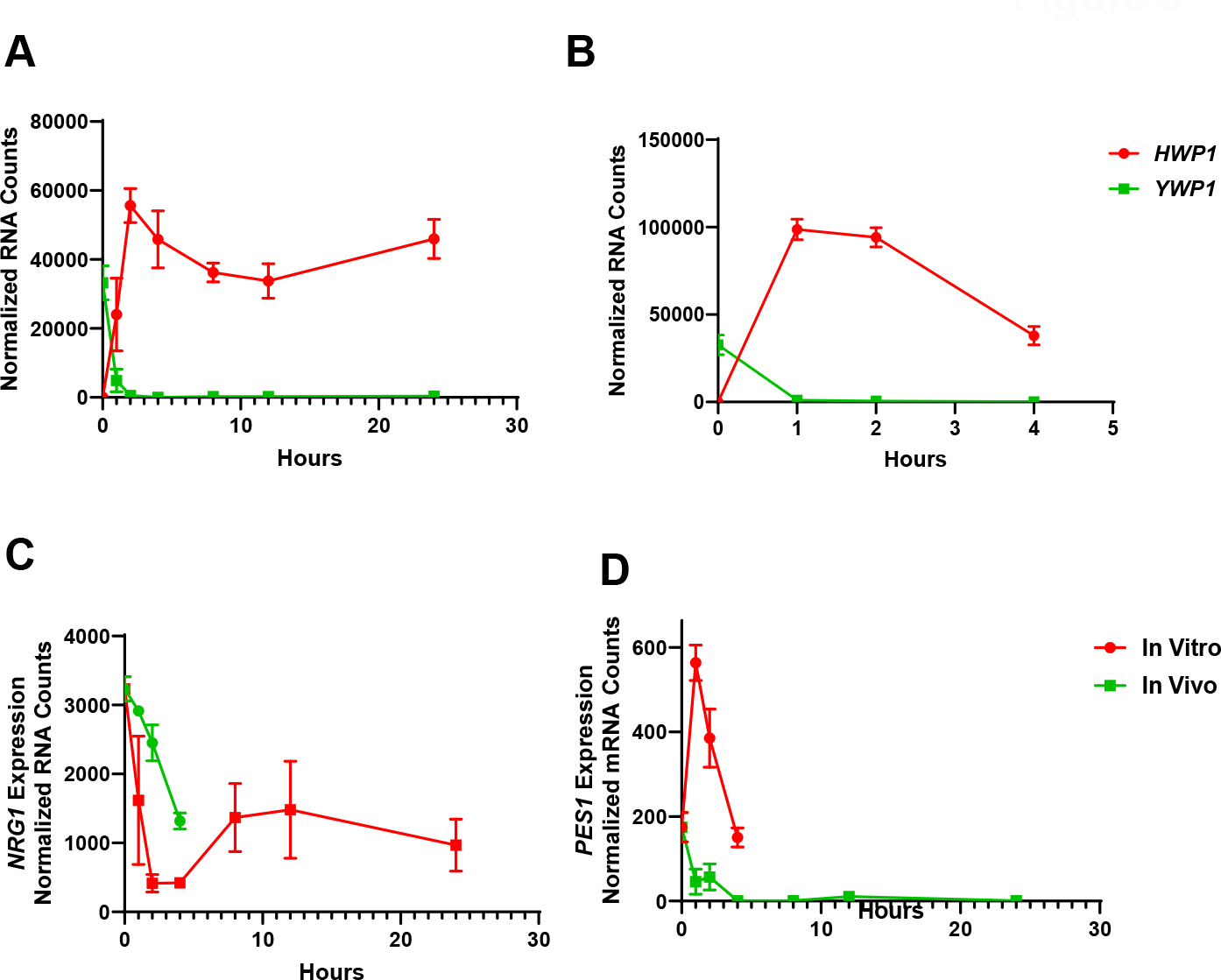
Time course of *HWP1, YWP1, NRG1*, and *PES1* expression during in vitro and in vivo filamentation. **A.** Normalized mRNA counts for *HWP1* and *YWP1* over time course of in vivo (**A**) and in vitro (**B**) filamentation (each point is mean of 3 mice or independent cultures; error bars are ± SD). Expression of *NRG1* (**C**) and *PES1* (**D**) over in vitro and in vivo time course.

*NRG1* is a repressor of hypha-associated genes (18, 19). During in vitro hyphal induction, its expression and protein levels are reduced (18, 20). We confirmed that pattern of *NRG1* expression in our in vitro conditions (Fig. 3C). In vivo, *NRG1* expression rapidly reaches a nadir (2hr) and then expression increases after 4 hr to a steady state that is maintained through the remainder of the time course. This biphasic pattern of expression is also observed in other induction media; specifically, *NRG1* mRNA and protein levels decrease initially and then return to near yeast levels of expression. Relative in vitro studies with YPD supplemented with BCS have shown NRG1 expression reaches a nadir at 1hr and is recovering by 2-3hr (21); in contrast, the recovery phase seems to be delayed in RPMI+10%BCS. Overall, these data indicate that *NRG1* dynamics are reasonably well conserved between in vivo and in vitro induction. Once again, the transcriptional steady state is reached prior to the establishment of a morphological steady state.

As noted above, one of the differences between the time courses of filamentation in vitro and vivo is the lack of lateral yeast formation in vivo. *PES1* is required for lateral yeast cell formation in vitro and is essential for yeast phase growth (15). In vitro, *PES1* expression is induced during filamentation relative to yeast phase, although it returns to yeast phase levels at 4hr (Fig. 3D); this same trend was observed in YPD+BCS (15). In contrast, *PES1* expression in vivo rapidly drops to near background levels and remains so throughout the time course (Fig. 3D). This observation provides an explanation for why lateral yeast are not observed during the 24hr time course of in vivo filamentation because *PES1* is required for the hyphae-to-yeast transition (15).

### The tempo of hyphae-specific gene and hyphal phase transcriptional factor expression suggests that in vivo cells maintain the hyphal program throughout the 24 hr time course

Under both in vitro and in vivo conditions, the expression of other canonical hyphae-associated genes is rapidly induced (Fig. 4A&B). In vivo, this a plateau is again observed. In vitro, the expression of all four genes is reduced at the 4 hr timepoint, consistent with the cells undergoing a hyphae-to-yeast transition. The expression of transcription factors that are induced during hyphal morphogenesis and are required for execution of its program also shows the rapid increase in expression and plateau in vivo (Fig. 4C) while in vitro there is a more pronounced decrease in expression after the peak at 1 hr (Fig. 4D). The reduction in *UME6* expression is particularly pronounced in vitro while in vivo the expression remains quite stable throughout the time course. The Kadosh lab has shown that maintenance of hyphal morphogenesis in vitro is modulated by the expression level of *UME6* (22). We have also showed that *UME6* is required for hyphal extension in vivo (7). The reduced expression of *UME6* and the other inducible positive transcriptional regulators of hypha formation at later time points in the in vitro experiment is consistent with the cells beginning to transition to the hyphae-to-yeast phase of morphogenesis. In contrast, the relatively stable expression of regulators such as *UME6* in vivo suggests that the cells are largely in the maintenance phase of filamentation in vivo.

**Figure 4.**
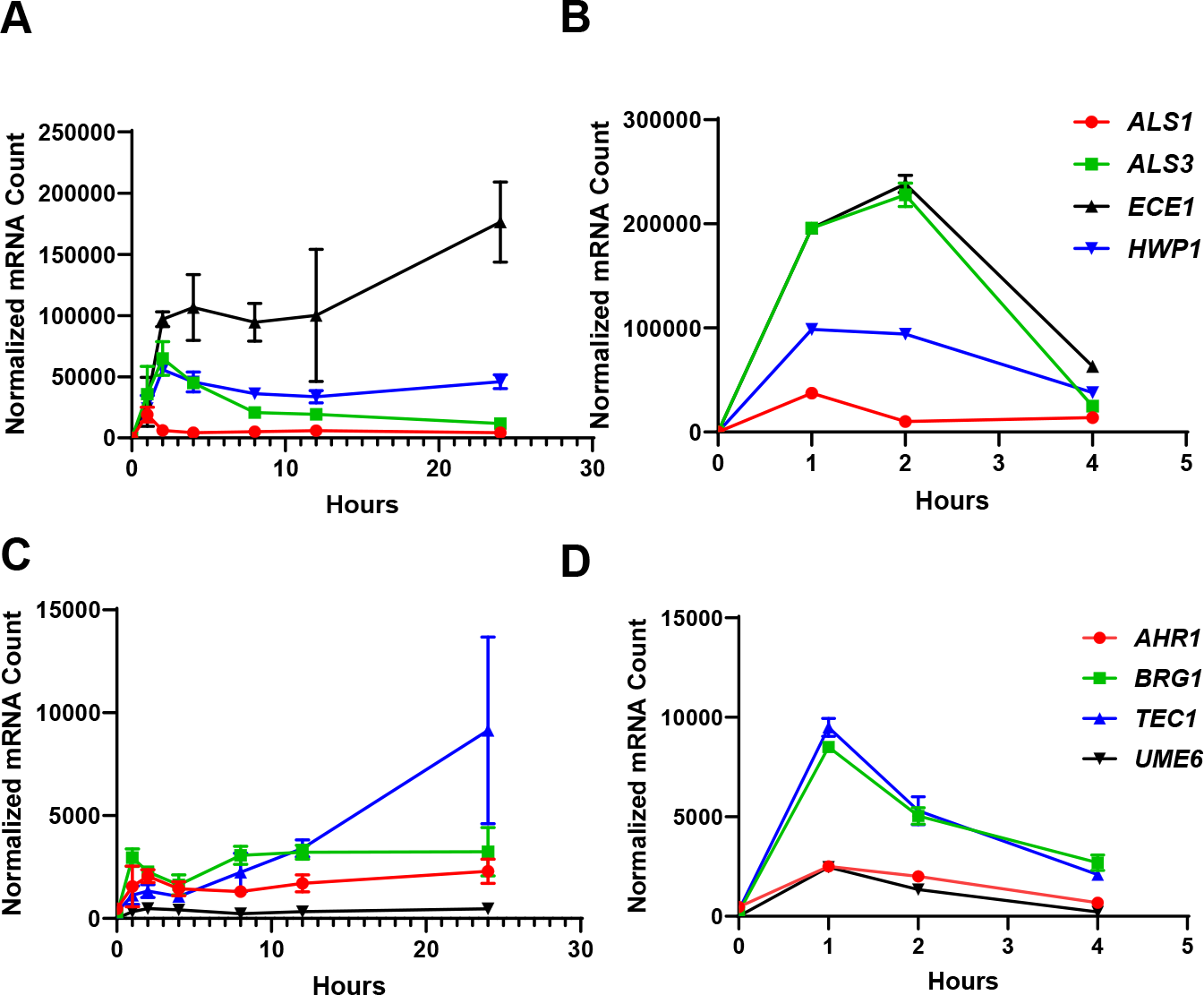
Time course of representative hyphae-specific genes and hyphae-associated transcription factors. Data are normalized counts of the indicated hyphae specific genes during in vivo (**A**) and in vitro (**B**) filamentation and indicated hyphae-associated transcription factors during in vivo (**C**) and in vitro (**D**) filamentation. Data points are mean of normalized mRNA counts from three mice or three independent cultures; error bars are ±SD.

### The transcriptional state of *C. albicans* filaments continues to remodel after initiation of hyphal program in vivo

The hyphal expression profile appears to be rapidly induced both in vivo and in vitro. To identify genes that are differentially expressed after the hyphal program is initiated, we normalized our in vivo data set to the 1 hr post-infection time point. While very few additional genes are upregulated after 2hr, the transcriptional profile continues to evolve over the remainder of the time course with 108 genes differentially expressed at 24hr. Indeed, 55 more genes are differentially expressed at 24hr relative to 12hr, the time point at which the ratio of filaments to yeast has reached steady state (Fig. 5A). The gene with the largest FC at 24hr relative to 1hr post-infection is *AOX2* which encodes one of two isoforms of the mitochondrial alternative oxidase (23). The expression of *AOX2* is below background until 8 hr but is upregulated 4000-fold by 24 hr (Fig. 5B). In vitro, *AOX2* is induced in the presence of poor carbon sources (23), suggesting that the metabolic state of the filaments is in flux during infection. Further supporting this conclusion, *PCK1*, a key gluconeogenic enzyme, is also induced at later time points (Fig. 5B). Finally, 6/8 of the ALS family of adhesion genes are downregulated at 24hr relative to 1hr post infection (Fig. 5C). In contrast, 5/8 PGA cell wall proteins are upregulated at the same time point (Fig. 5D). This suggests that the cell wall/cell surface proteins of the filaments are also under remodeling after initial hyphal program induction.

**Figure 5.**
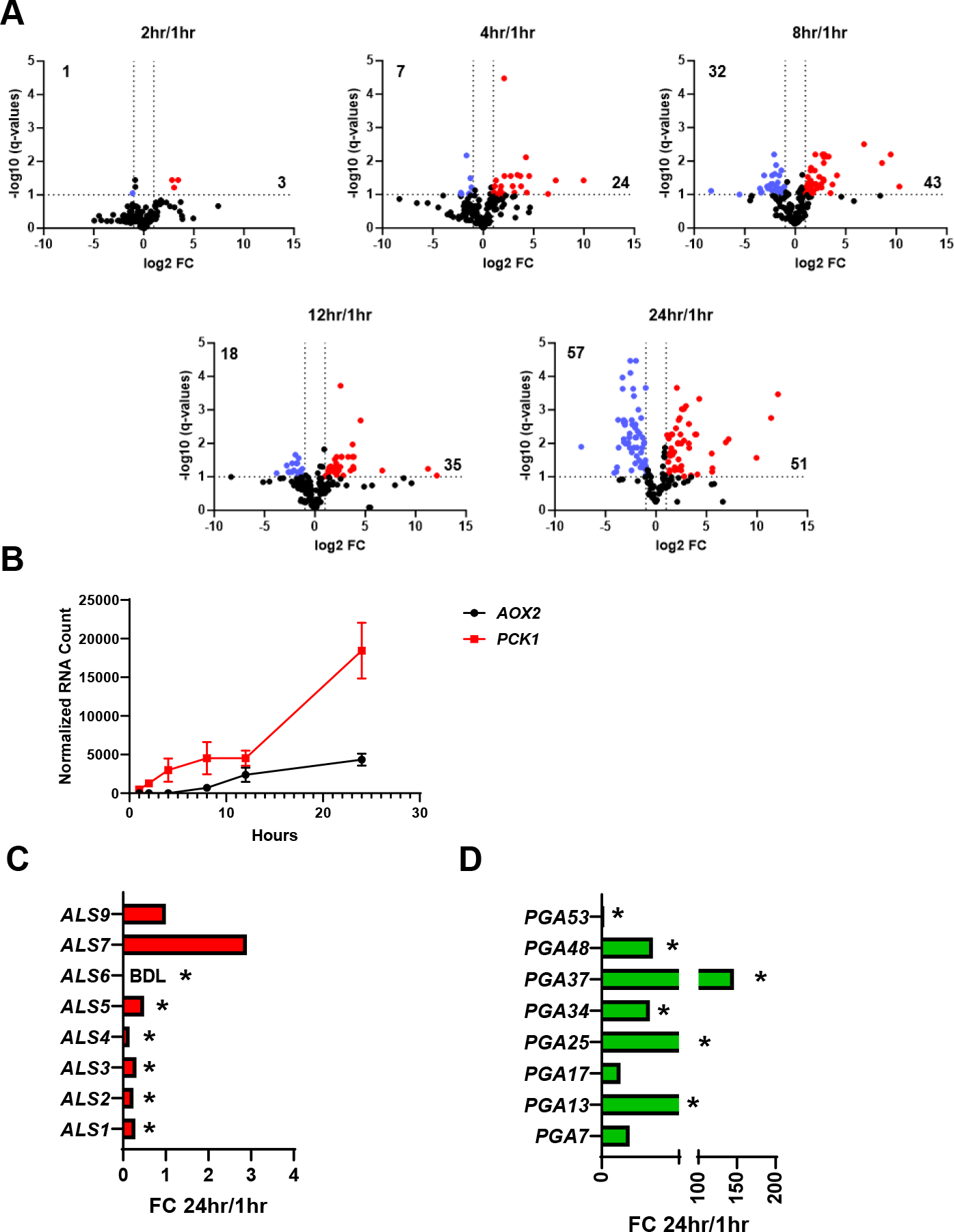
The transcriptional profile during in vivo filamentation remains dynamic after initial induction of hyphae-associated genes. **A.** The data depicted in Fig. 2 were renormalized to 1 hr post-infection to control for induction of the hyphal program. Volcano plots showing genes with significant (log_2_ ± 1; FDR<0.1, Benjamini method) increase (red dots) or decrease (blue dots) at the indicated time points. The numbers in the two quadrants indicate total number of differentially expressed genes. **B**. Expression of *AOX2* and *PCK1* over the time course of in vivo filamentation. Data points are mean of normalized mRNA counts from three mice; error bars are ±SD. Fold change in ALS family genes (C) and PGA family genes (D) at 24hr relative to 1 hr. Asterisk indicate that the difference in gene expression is statistically significant (FDR<0.1, Benjamini method).

## Discussion

Here, we report the first temporal analysis of *C. albicans* gene expression during in vivo filamentation and a comparison those dynamics with in vitro filamentation under widely used “host-like” conditions. Our data indicate that initiation of the hyphal gene expression program occurs rapidly under both in vitro and in vivo conditions with the induction of hyphae-associated genes and hyphal transcriptional regulators occurring within 1-2 hr. Under both conditions, the expression of these hyphae-associated genes reaches a maximum level prior to a morphological steady state is reached. In vitro, the expression of the hyphae-associated genes begins to decline by the end of the 4 hr induction period. Microarray-based time course analysis of in vitro filamentation reported by Kadosh and Johnson (12) showed a similar decrease in expression of hyphae-associated genes in different medium (YPD+10% BCS, 37°C). We suggest that this decline in hyphal gene and transcription factor expression is related to a transition toward the hyphae-to-yeast transition that begins to occur at ∼4hr and increases thereafter in vitro (14, 15). Consistent with that notion is the observation, that *PES1* expression actually increases in vitro and is at yeast phase expression at 4hr. Pes1 is required for lateral yeast formation and overexpression of *PES1* drives lateral yeast formation in vitro (15).

In vivo, the increased expression of hyphae-associated genes remains at a plateau throughout the 24 hr time course and after a morphological steady state has been reached. Transcription factors that are required for WT levels of filamentation and filament length such as *TEC1, BRG1*, and *UME6* are expressed near or in excess of the levels established after 1-2hr post-infection, indicating that the hyphal transcriptional program remains operative throughout the time course. Further supporting that conclusion, the expression of *PES1* is below our detection limit after the 2 hr time point. As noted above, this observation provides an explanation for why lateral yeast formation is not observed in vivo over this time course. Taken together, these data indicate that in vitro filamentation under the conditions we studied occurs over a much more compressed time scale compared to the in vivo conditions of the ear model despite the initiation of filamentation occurring with similar dynamics.

We also observed that hyphae remain transcriptionally dynamic after induction of the hyphal program in vivo. For example, the expression of the alternate oxidase *AOX2* is initially below detection but then is highly expressed at later timepoints. Aox2 reduces molecular oxygen to water on the inner mitochondrial membrane using ubiquinol as the electron source and without pumping electrons across the membrane (23). Aox2 is the inducible isoform of alternative oxidase in *C. albicans* while Aox1 is constitutively expressed. Recently, Liu et al. showed that *AOX2* is required for full virulence in a disseminated model of candidiasis but is dispensable for colonization of the GI tract (23). Consistent with that finding, Nanostring analysis of the expression profile of *C. albicans* in the GI tract relative to yeast cells in YPD (the same conditions as our in vitro yeast reference) showed that *AOX2* is downregulated dramatically in all segments of the intestine (4). Our data show that *AOX2* is downregulated initially relative to the YPD-grown inoculum but then increases in expression after morphological steady-state is reached. Histological analysis of kidneys infected with the *aox2*ΔΔ mutant indicated that hyphal elements were present (23), suggesting that its virulence defect is independent of morphogenesis in vivo.

Our data suggest that *AOX2* expression does not occur until after hyphae have been formed in vivo and, therefore, provides a potential explanation for why the *aox2*ΔΔ mutant both forms filaments and is less virulent.

In vitro, *AOX2* is induced when cells are cultivated in poorly fermentable or non-fermentable carbon sources such as galactose and glycerol. This suggests that, as the infection proceeds, glucose becomes less abundant, triggering the filaments to shift their metabolism to other available carbon sources. Further supporting this change in carbon metabolism, the key gluconeogenic enzyme *PCK1* is also induced as the infection proceeds. *PCK1* expression is suppressed by glucose in vitro and induced by lactate and succinate (24). *PCK1* is also downregulated in the GI tract (4).

The ear infection model is based on infecting the subepithelial tissue that is anatomically similar to stromal tissue beneath epithelial and mucosal tissue (6, 7, 8). A current model for pathogenesis of invasive candidiasis involves translocation of GI-resident *C. albicans* across the gut mucosae to invade the submucosal tissue. This process is thought to be dependent upon filamentation. Our data regarding the expression of *AOX2* and *PCK1* during filamentation in this compartment suggests that the carbon sources available after filamentation in the submucosae require a significant shift in metabolism compared to cells commensally resident in the GI tract. *SAP6* is a gene whose expression reduces fitness in the GI tract (4) and is upregulated strongly during filamentation (442-fold relative to yeast, Table S1) in the subepithelial compartment of the ear model. Conversely, *CHT2* is a gene that is upregulated during, and promotes, commensalism in the GI tract and is downregulated during in vivo filamentation (Table S1) in the subepithelial stroma (25). Thus, dynamics of *C. albicans* gene expression during in vivo filamentation supports developing models for the distinctions between commensal and invasive phases of *C. albicans* pathophysiology.

It is important to discuss the limitations of this analysis. First, we have focused on a set of specific time points for our studies; it is likely that extension of the analysis to later timepoints would lead to alternate or additional findings. We have also used a single in vitro condition as a comparator for the in vivo experiments. Second, many in vitro conditions can be used to induce filamentation and, as we and others have shown, the specific conditions used to study *C. albicans* has a significant effect on gene expression and the phenotypes of some mutants (8, 13). Third, this potential limitation also applies to the specific site or organs that are studied in in vivo which are also likely to have distinct gene expression profiles during filamentation (9, 26). Fourth, our studies have focused on a specific set of genes related to hyphae formation and adaptation to the environment. It is likely that other genes are also undergoing changes. Accordingly, we have focused our analysis of genes that act as physiologic reporters of well-defined cell states or physiologies.

Despite these limitations, we can conclude with confidence that: 1) transcriptional programs regulating filamentation are rapidly induced in vitro and in vivo; 2) the tempo of filamentation in vivo is prolonged relative to in vitro filamentation and the period of high expression of genes associated with that process is also prolonged; 3) hyphae are adapting to changing infection environments after filamentation has reached steady-state; and 4) the expression of key genes involved in virulence and commensalism are consistent with genetic evidence supporting their roles in those two processes.

These data also contribute to the developing concept that *C. albicans* filamentation is intricately contingent on the specific environment with in which it occurs. As such, it is important to regard the filamentation process as a separate growth phase of *C. albicans* that is as adaptable to changing growth conditions as the more familiar yeast phase. From this perspective, the hyphal state is not special case defined only by a canonical set of genes whose expression mark that state but rather a dynamic morphology that is responsive to the environment and the host.

## Materials and methods

### Strains and media

The *C. albicans* reference strain SN250 was used for all experiments. All *C. albicans* strains were precultured overnight in yeast peptone dextrose (YPD) medium at 30°C with shaking. Standard recipes were used to prepare YPD (27). RPMI 1640 medium was supplemented with bovine calf serum (10% vol/vol).

### In vitro hyphal induction

For in vitro hyphal induction, *C. albicans* strain was incubated overnight at 30°C in YPD media, harvested, and diluted into RPMI + 10% bovine calf serum at a 1:50 ratio and incubated at 37°C. Cells were collected at the different time points (e.g., 1hr, 2hr, and 4hr) and processed for microscopy or RNA isolation as described below.

### In vitro characterization of *C. albicans* morphology

Induced cells were fixed with 1% (v/v) formaldehyde. Fixed cells were then imaged using the Echo Rebel upright microscope with a 60x objective. The assays were conducted in triplicates on different days to confirm reproducibility.

### In vivo characterization of *C. albicans* morphology

These assays were performed as previously described (6). Briefly, 1 X 10^6^ WT *C. albicans* cells were inoculated intradermally in mouse ear (3 mice/time point). Mice (3 mice/time point) were sacrificed at each time points, and ears were harvested. One ear/mouse was immediately submerged into the ice-cold RNA later solution and another ear used for the imaging. A multiple Z stacks (minimum 20) were acquired and used it to score the yeast vs filamentous ratio. Round and/or budded cells were considered “Yeast”, whereas if the cells contain intact mother and filamentous which was at least twice the length of the mother body, were considered “filamentous.” A minimum of 100 cells from multiple fields were scored. Student’s *t* test was performed to define the statistical significance between the different time points. All animal experiments were approved by the University of Iowa IACUC.

### RNA extraction

In vitro and in vivo RNA extraction was carried out as described previously (7, 9). For in vitro RNA extraction, cells were collected at the different time points, centrifuged for 2 min at 10K rpm at room temperature and RNA was extracted according to the manufacturer protocol (MasterPure Yeast RNA Purification Kit). For in vivo RNA extraction, mouse was euthanized following the protocol approved by the University of Iowa IACUC. Mouse ear was cut carefully and placed into the ice-cold RNA later solution. The ear was then transferred to the mortar and flash frozen with liquid nitrogen. Using pestle, the frozen was ground to the fine powder. The resulting powder was collected and 1 ml of ice-cold Trizol was added. The samples were placed on a rocker at RT for 15 min and then centrifuged at 10K at 4° C for 10 min. The cleared Trizol was collected into 1.5 ml Eppendorf tube and 200 μl of RNase free chloroform was added to each sample. The tubes were shaken vigorously for 15 s and kept at RT for 5 min followed by centrifuge at 12K rpm at 4° C for 15 min. The cleared aqueous layer was then collected to a new 1.5 ml Eppendorf tube and RNA was further extracted following the Qiagen RNeasy kit protocol.

### NanoString® gene expression analysis

NanoString analysis was carried out as described previously (7, 9). Briefly, in total, 40 ng of *in vitro* or 1.4 μg of *in vivo* RNA was added to a NanoString codeset mix and incubated at 65° C for 18 hours. After hybridization reaction, samples were proceeded to nCounter prep station and samples were scanned on an nCounter digital analyzer. nCounter .RCC files for each sample were imported into nSolver software to evaluate the quality control metrics. Background subtraction was performed using negative control probes to establish a background threshold, which was then subtracted from the raw counts. The resulting background subtracted total raw RNA counts underwent a two-step normalization process. First normalized against the highest total counts from the biological triplicates and then to the wild type samples. The statistical significance of changes in gene expression was determined using Benjamini-Hochberg method at an FDR of 0.1.

### Software

All the image analysis was carried out using ImageJ software. Graph pad prism (V. 9.3.1) was used to plot the graphs and to perform the statistical tests.

## Acknowledgements

The authors thank Scott Filler (UCLA) for helpful discussions and critical review of the manuscript. This work was supported by a National Institutes of Health Grant, R01AI133409 (DJK).

## Supplementary Tables

**Table S1.** Nanostring data for in vivo time course with raw counts, background corrected counts, normalized counts, fold change for each gene relative to either yeast phase or 1hr postinfection, t-test values, and adjusted q values corresponding to FDR calculated by the Benjamini method.

**Table S2.** Nanostring data for in vitro time course with raw counts, background corrected counts, normalized counts, fold change for each gene relative to yeast phase, t-test values, and adjusted q values corresponding to FDR calculated by the Benjamini method.

